# Geometry of the sample frequency spectrum and the perils of demographic inference

**DOI:** 10.1101/233908

**Authors:** Zvi Rosen, Anand Bhaskar, Sebastien Roch, Yun S. Song

## Abstract

The sample frequency spectrum (SFS), which describes the distribution of mutant alleles in a sample of DNA sequences, is a widely used summary statistic in population genetics. The expected SFS has a strong dependence on the historical population demography and this property is exploited by popular statistical methods to infer complex demographic histories from DNA sequence data. Most, if not all, of these inference methods exhibit pathological behavior, however. Specifically, they often display runaway behavior in optimization, where the inferred population sizes and epoch durations can degenerate to 0 or diverge to infinity, and show undesirable sensitivity of the inferred demography to perturbations in the data. The goal of this paper is to provide theoretical insights into why such problems arise. To this end, we characterize the geometry of the expected SFS for piecewise-constant demographic histories and use our results to show that the aforementioned pathological behavior of popular inference methods is intrinsic to the geometry of the expected SFS. We provide explicit descriptions and visualizations for a toy model with sample size 4, and generalize our intuition to arbitrary sample sizes *n* using tools from convex and algebraic geometry. We also develop a universal characterization result which shows that the expected SFS of a sample of size n under an *arbitrary* population history can be recapitulated by a piecewise-constant demography with only *κ_n_* epochs, where *κ_n_* is between *n*/2 and 2*n* – 1. The set of expected SFS for piecewise-constant demographies with fewer than *κ_n_* epochs is open and non-convex, which causes the above phenomena for inference from data.

## 1 Introduction

The sample frequency spectrum (SFS), also known as the site or allele frequency spectrum, is a fundamental statistic in population genomics for summarizing the genetic variation in a sample of DNA sequences. Given a sample of *n* sequences from a panmictic (i.e., randomly mating) population, the SFS is a vector of length *n* – 1 of which the *k*th entry corresponds to the number of segregating sites each with *k* mutant (or derived) alleles and *n* – *k* ancestral alleles. The SFS provides a compact way to summarize *n* sequences of arbitrary length into just *n* – 1 numbers, and is frequently used in empirical population genetic studies to test for deviations from equilibrium models of evolution. For instance, the SFS has been widely used to infer demographic history where the effective population size has changed over time (Nielsen, 2000; Gutenkunst et al., 2009; Gravel et al., 2011; Keinan and Clark, 2012; Excoffier et al., 2013; Bhaskar et al., 2015), and to test for selective neutrality (Kaplan et al., 1989; Achaz, 2009). In fact, many commonly used population genetic statistics for testing neutrality, such as Watterson’s *θ_W_* (Watterson, 1975), Tajima’s *θ_π_* (Tajima, 1983), and Fu and Li’s *θ_FL_* (Fu and Li, 1993) can be expressed as linear functions of the SFS (Durrett, 2008).

In the coalescent framework (Kingman, 1982a,b,c), the *unnormalized expected* SFS **ξ**_*n*_ for a random sample of *n* genomes drawn from a population is obtained by taking the expectation of the SFS over the distribution of sample genealogical histories under a specified population demography. In this work, we will be concerned with well-mixed, panmictic populations with time-varying historical population sizes, evolving according to the neutral coalescent process with the infinite-sites model of mutation. The coalescent arises as the continuum limit of a large class of discrete models of random mating, such as the Wright-Fisher, Moran, and Cannings exchangeable family of models (Möhle and Sagitov, 2001). The infinite-sites model postulates that every mutation in the genealogy of a sample occurs at a distinct site, and is commonly employed in population genetic studies for organisms with low population-scaled mutation rates, such as humans. The SFS also appears in the context of statistical modeling as a vector of probabilities. In particular, the *normalized expected* SFS 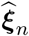, defined by normalizing the entries of **ξ**_*n*_ so that they sum to 1, gives the probability that a random mutation appears in *k* out of n sequences in the sample. Unless stated otherwise, we use the term expected SFS to refer to the unnormalized quantity **ξ**_*n*_.

The expected SFS is strongly influenced by the demographic history of the population, and extensive theoretical and empirical work has been done to characterize this dependence (Fu, 1995; Wakeley and Hey, 1997; Polanski et al., 2003; Marth et al., 2004; Chen, 2012; Kamm et al., 2017; Jouganous et al., 2017). Fu (1995) showed that under the infinite-sites model for a panmictic population with constant size and no selection, the expected SFS is given by 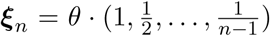, where θ/2 denotes the population-scaled mutation rate. When the population size is variable, however, the formula for the expected SFS depends on the entire population size history. In particular, Polanski and Kimmel (2003) showed that the expected SFS under a time-varying population size is given by **ξ**_*n*_ = *A_n_c*, with *A_n_* being an (*n* – 1)-by-(*n* – 1) invertible matrix that only depends on *n* and *c* = (*c*_2_,…, *c_n_*), where *c_m_* denotes the expected time to the first coalescence event in a random sample of size *m* drawn from the population at present. For any time-varying population size function *η*(*t*), *c_m_* is given by the following expression:

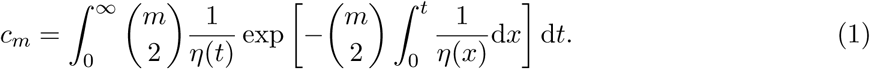

A natural statistical question that arises when using the SFS for demographic inference is whether it is theoretically possible to reconstruct the population history *η*(*t*) from the expected SFS **ξ**_*n*_ of a large enough sample size *n*. This question was famously answered by Myers et al. (2008) in the negative, by constructing a way of perturbing any population size history without altering the expected SFS for all sample sizes. However, in subsequent work by two of this paper’s authors (Bhaskar and Song, 2014), it was shown that for a wide class of biologically plausible population histories, such as those given by piecewise-constant and piecewise-exponential functions, the expected SFS of a finite sample size is sufficient to uniquely identify the population history. These two results give us insight into the map from population size histories to the expected SFS vectors. The space of *all* possible population size histories is of infinite dimension, while the expected SFS vectors for any fixed sample size *n* form a finite-dimensional space; naturally, the pre-image of an expected SFS under this map will typically be an infinite set of population size histories. However, imposing conditions such as those of Bhaskar and Song (2014) restricts us to a function space of finite dimension, so that the pre-images of the expected SFS may become finite/unique.

While the above results are concerned with the identifiability of demographic models from noiseless SFS data, they do not directly provide an explicit characterization of the geometry of the expected SFS as a function of the demographic model. Studying such geometry would be very useful for understanding the behavior of inference algorithms which perform optimization by repeatedly computing the image of the map from the space of demographic parameters to the expected SFS while trying to minimize the deviation of the expected SFS from the observed SFS data. To this end, our main contribution is a universal characterization of the space of expected SFS for any sample size n under *arbitrary population size histories*, in terms of the space of expected SFS of piecewise-constant population size functions with *O*(*n*) epochs. This is a useful reduction because the latter space is much more tractable for mathematical analysis and computation. We provide a complete geometric description of this space for a sample of size *n* = 4, and generalize our intuition to arbitrary sample sizes *n* using tools from convex and algebraic geometry. Our reduction also provides an explanation for a puzzling phenomenon frequently observed in empirical demographic inference studies - namely, for some observed SFS data, the optimization procedure for inferring population histories sometimes exhibits pathological behavior where the inferred population sizes and epoch durations can degenerate to 0 or diverge to ∞.

## 2 Piecewise-Constant Demographies

Let Π_*k*_ be the set of piecewise-constant population size functions with *k* pieces. Any population size function in Π_*k*_ is described by 2*k* – 1 positive numbers, representing the *k* population sizes (*y*_1_,…, *y_k_*) and the *k* – 1 time points (*t*_1_,…, *t*_*k*-1_) when the population size changes. Let *Ξ_n,k_*, which we call the (*n,k*)-SFS manifold^*^, denote the set of all expected SFS vectors for a sample of size *n* that can be generated by population size functions in Π_*k*_. Similarly, let *𝒞_n,k_*, called the (*n, k*)-coalescence manifold, denote the set of all vectors c = (*c*_2_,…, *c_n_*) giving the expected first coalescence times of samples of size 2,…,*n* for population size functions in Π_*k*_. Let 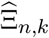 and 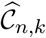 respectively be equal to the normalization of all points in *Ξ_n,k_* and *𝒞_n,k_* by their *l*_1_-norms (i.e., the sums of their coordinates). Note that both manifolds live in ℝ^*n*−1^ and their normalized versions live in the (*n* – 2)-dimensional simplex Δ^*n*−2^; this is the set of nonnegative vectors in ℝ^*n*−1^ whose coordinates sum to 1.

Now that we have defined our basic objects of study, we can describe the remainder of the paper: In Section 3, we provide a complete geometric picture of the Ξ_4,*k*_ SFS manifold describing the expected SFS for samples of size *n* = 4 under piecewise-constant population size functions with an arbitrary number *k* of pieces. We make explicit the map between regions of the demographic model space and the corresponding probability vectors, and this will foreshadow some of the difficulties with population size inference in practice. In Section 4, we develop a characterization of the space of expected SFS for arbitrary population size histories. In particular, we show that for any sample size n, there is a finite integer *κ_n_* such that the expected SFS for a sample of n under any population size history can be generated by a piecewise-constant population size function with at most *κ_n_* epochs. Stated another way, we show that the Ξ_*nκ_n_*_ SFS manifold contains the expected SFS for all possible population size histories, no matter how complicated their functional forms. We establish bounds on *κ_n_* that are linear in *n*, and along the way prove some interesting results regarding the geometry of the general Ξ_*n,k*_, SFS manifold. Finally, in Section 5, we demonstrate the implications of our geometric characterization of the expected SFS for the problem of demographic inference from noisy genomic sequence data.

Before proceeding further, we state a proposition regarding the structure of the map from Π_*k*_ to *𝒞_n,k_*, which we will call 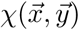 the vector of *k* – 1 transformed breakpoints is denoted by 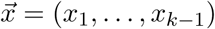 and defined below, while the vector of population sizes in the *k* epochs is denoted by 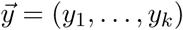. This allows us to explain the algebraic nature of most of our proofs. All proofs of the results presented in this paper are deferred to Section 7.

### Proposition 2.1

*Fix a piecewise-constant population size function in* Π_*k*_ *with epochs* [*t*_0_, *t*_1_), [*t*_1_,*t*_2_),…, [*t*_*k*-1_,*t_k_*), where 0 = *t*_0_ <*t*_1_ < … < *t*_*k*-1_ <*t_k_* = ∞, *and which has constant population size value y_j_ in the epoch* [*t*_*j*-1_, *t_j_*) *for j* = 1,…,*k. Let x_j_* = exp[–(*t_j_ – *t*_*j*−1_)/y_j_* ] *for j* = 1,…, *k, where x_k_* = 0 (corresponding to time T = ∞), *and define x*_0_ = 1 (*corresponding to time T* = 0) *for convenience. The vectors* (*x*_1_,…, *x*_*k*-1_,*y*_1_,…,*y_k_*), *where* 0 < *x_j_* < 1 *and y_j_* > 0 *for all j, (uniquely) identify the population size functions in* Π_*k*_, *and they satisfy both of the following equations*:

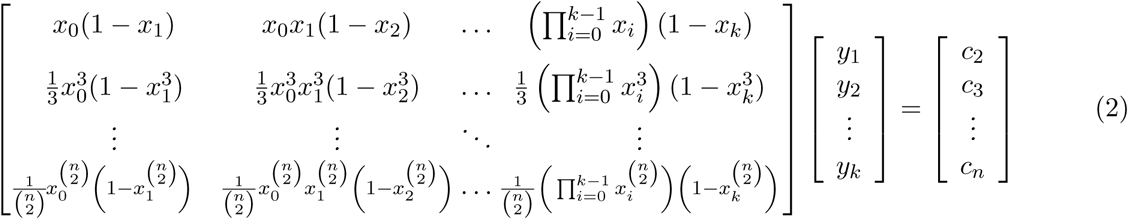

and

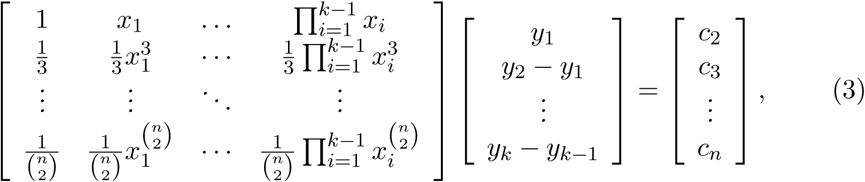

*where c_m_ is the expected first coalescence time for a sample of size m, as defined in* (1).

These two formulations provide two perspectives on the coalescence manifold *𝒞_n,k_*:

1. In (2), the left-hand matrix, call it *M*_1_(*n,k*), has each column of the same form with two parameters; this indicates they all live in a 2-dimensional surface. The vector (*y*_1_,…, *y_k_*) has all positive parameters. This means that the vector c = (*c*_2_,…, *c_n_* is contained in the cone over the surface described by the columns of *M*_1_.
2. In (3), the left-hand matrix, call it *M*_2_(*n,k*) has each column of the same form with one parameter; this indicates they all live on a curve. The vector (*y*_1_,*y*_2_ – *y*_1_, …,*y_k_* – *y*_*k*-1_) on the left hand side has parameters with possibly negative coordinates. So the vector c = (*c*_2_,…, *c_n_* is contained in the linear span of the curve described by the columns of *M*_2_.

Proposition 2.1 gives us the algebraic mappings that will serve as our objects of interest. Since the SFS manifold is simply a linear transformation of the coalescence manifold, we will use these maps as our entry into understanding the SFS manifold.

## 3 The Ξ_4,*k*_ SFS Manifold: A Toy Model

The first in-depth study will involve the set of all possible expected SFS for a sample of size 4. We choose *n* = 4 for a number of reasons: First, the cases of *n* = 2 and 3 are cones with simple boundaries in the line or plane. Second, when *n* = 4, the absolute SFS manifold lives in ℝ^3^, which can be nicely visualized, and the normalized SFS manifold lives in the 2-simplex, i.e. the triangle with vertices (1,0,0), (0,1,0), and (0,0,1). Finally, as observed in Proposition 2.1, the most interesting phenomena in SFS manifolds of any dimension are fundamentally phenomena of curves and surfaces. These are already captured in the *n* = 4 case.

For the sake of completeness, we begin by formally describing the coalescence manifolds *𝒞_n,k_* for the trivial cases of *n* = 2 and *n* = 3.

### Proposition 3.1

*We list some basic results on the coalescence manifolds 𝒞_n,k_ for small values of (n, k)*:

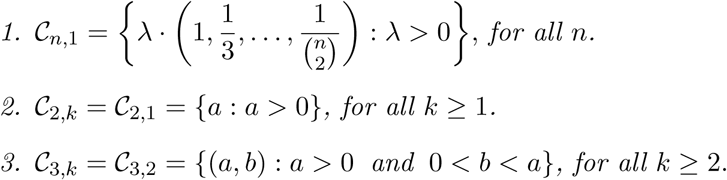

Note that from (2) and (3) for 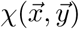 (Section 2), it follows that 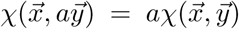 for *a* > 0. In words, rescaling the population sizes in each epoch by a constant *a* also rescales the first coalescence times by *a*. This implies that every point in the coalescence manifold *𝒞_n,k_* generates a full ray contained in the *𝒞_n,k_* coalescence manifold. Another consequence is that the normalized coalescence manifold 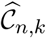 is precisely the intersection of the coalescence manifold *𝒞_n,k_* with the simplex Δ^*n*−2^.

With that justification, we begin to consider the normalized coalescence manifold 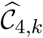 living in the simplex. As stated in Proposition 3.1, *𝒞*_4,1_ is a ray, which implies that 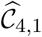 is a single point. We now characterize the set 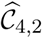.

### Proposition 3.2

*The manifold 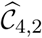 is a two-dimensional subset of the 2-simplex which can be described as the union of the point 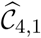 with the interiors of the convex hulls of two curves γ*_1_ *and γ*_2_. *The curves are parametrized as follows*:

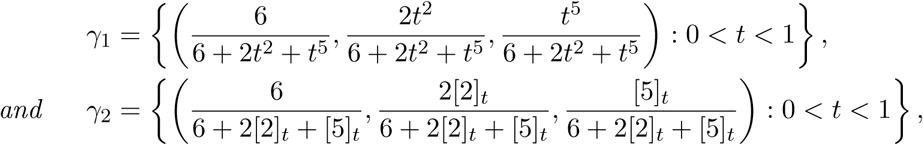

*where* [*n*]*_t_ denotes* 1 + … + *t^n^*.

This set has some highly unpleasant geometry. First of all, the set is non-convex; it is also neither closed nor open, because most of the boundary is excluded with the exception of the point (2/3; 2/9; 1/9). The set is visualized in Figure 1(a).

**Figure 1:**
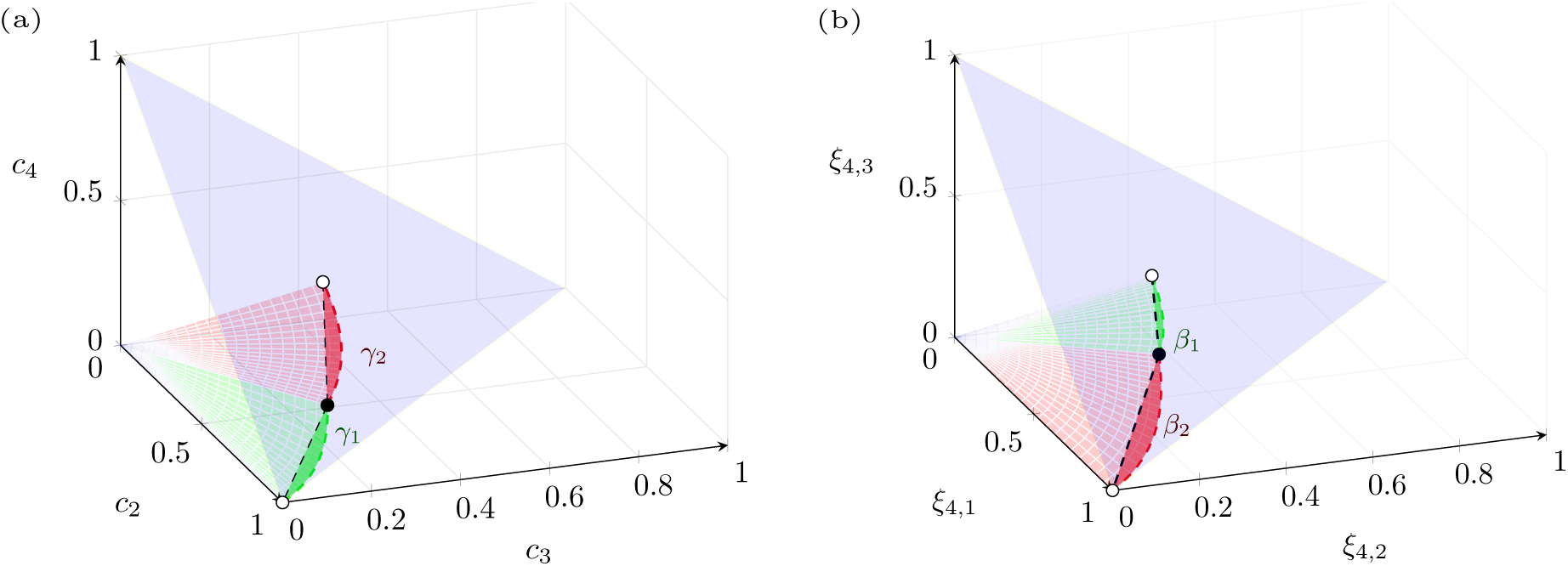
Coalescence and SFS manifolds for sample size 4 and 2 population epochs. (a) The coalescence manifold 𝓒_4,2_ is the union of red and green cones. The 2-simplex, shaded in blue, intersects 𝓒_4,2_ in the normalized coalescence manifold 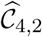. The green region corresponds to small-then-large demographies; the red region to large-then-small demographies. (b) The SFS manifold Ξ_4,2_ is the union of red and green cones. The 2-simplex intersects Ξ_4,2_ in the normalized SFS manifold 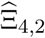. Here, too, the green region corresponds to small-then-large demographies; the red region to large-then-small demographies. Ξ_4,2_ is obtained from 𝓒_4,2_ by a linear transformation.

In order to precisely illustrate the geometry of 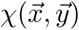 we will consider how contours in the domain map to contours in the image. Specifically, we plot the images of lines with fixed values of x_1_, respectively fixed values of (*y*_1_, *y*_2_), to 𝒞_4,2_ in the 2-simplex. The resulting contours are pictured in Figure 2.

**Figure 2:**
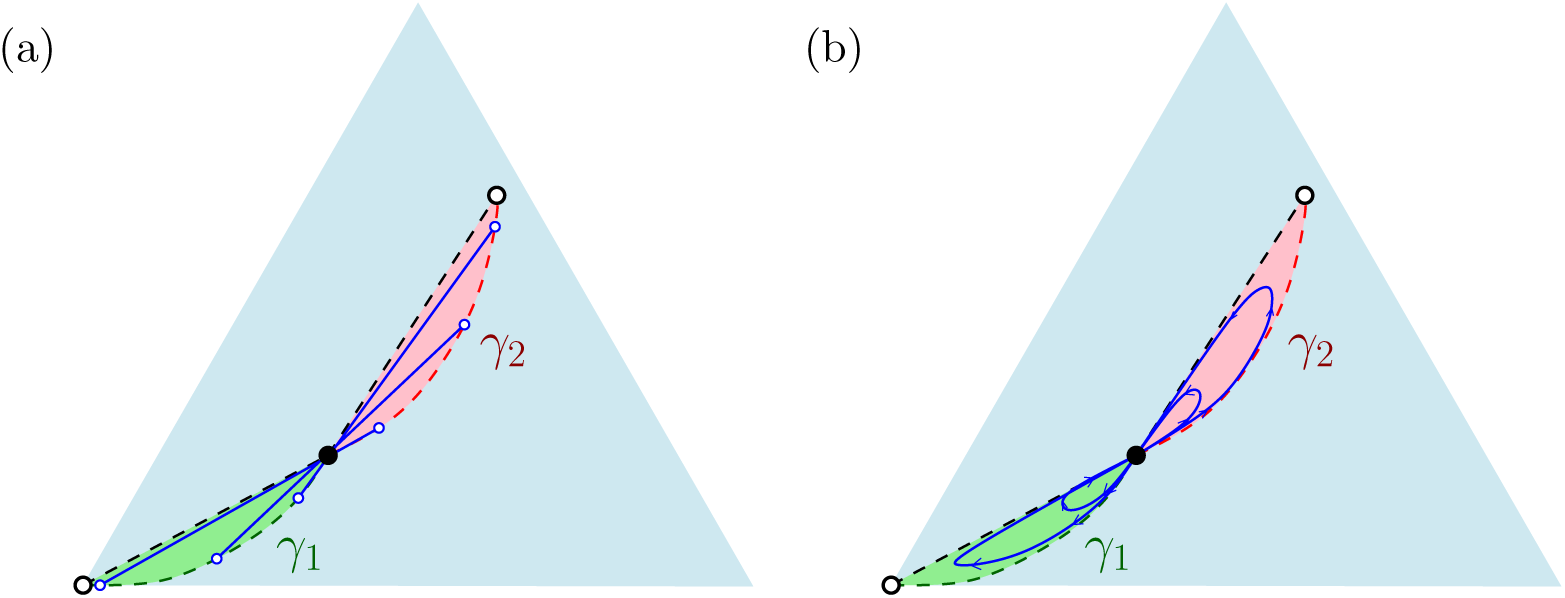
Fixed-time and fixed-size contours in 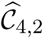. (a) The blue line segments correspond to the image of 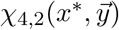 where *x*^*^ is a constant fixing the break-point between the two demographies. The other input 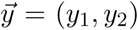 varies over all positive vectors, though scaled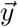 vectors point to the same normalized value. As *y*_1_/*y*_2_ → 0, the image approaches γ_2_ and as *y*_2_/*y*_1_→ 0, the image approaches γ_1_. (b) The blue curves correspond to the image of 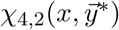 where 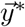 is a fixed vector indicating the population values and *x* takes all values in (0,1). The endpoints 0 and 1 correspond to breakpoints at ∞ and 0 respectively. For 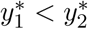, *x* traces a loop in the green region; for 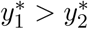, *x* traces a loop in the red region.

Finally, we consider how the map χ acts on the boundaries of the domain. To aid visualization, we limit the inputs to *x*_1_ and *y*_1_/*y*_2_, since all rescalings of *y*_1_ and *y*_2_ by the same positive constant while keeping *x*_1_ fixed map to the same normalized coalescence vector. The resulting map is illustrated in Figure 3.

**Figure 3:**
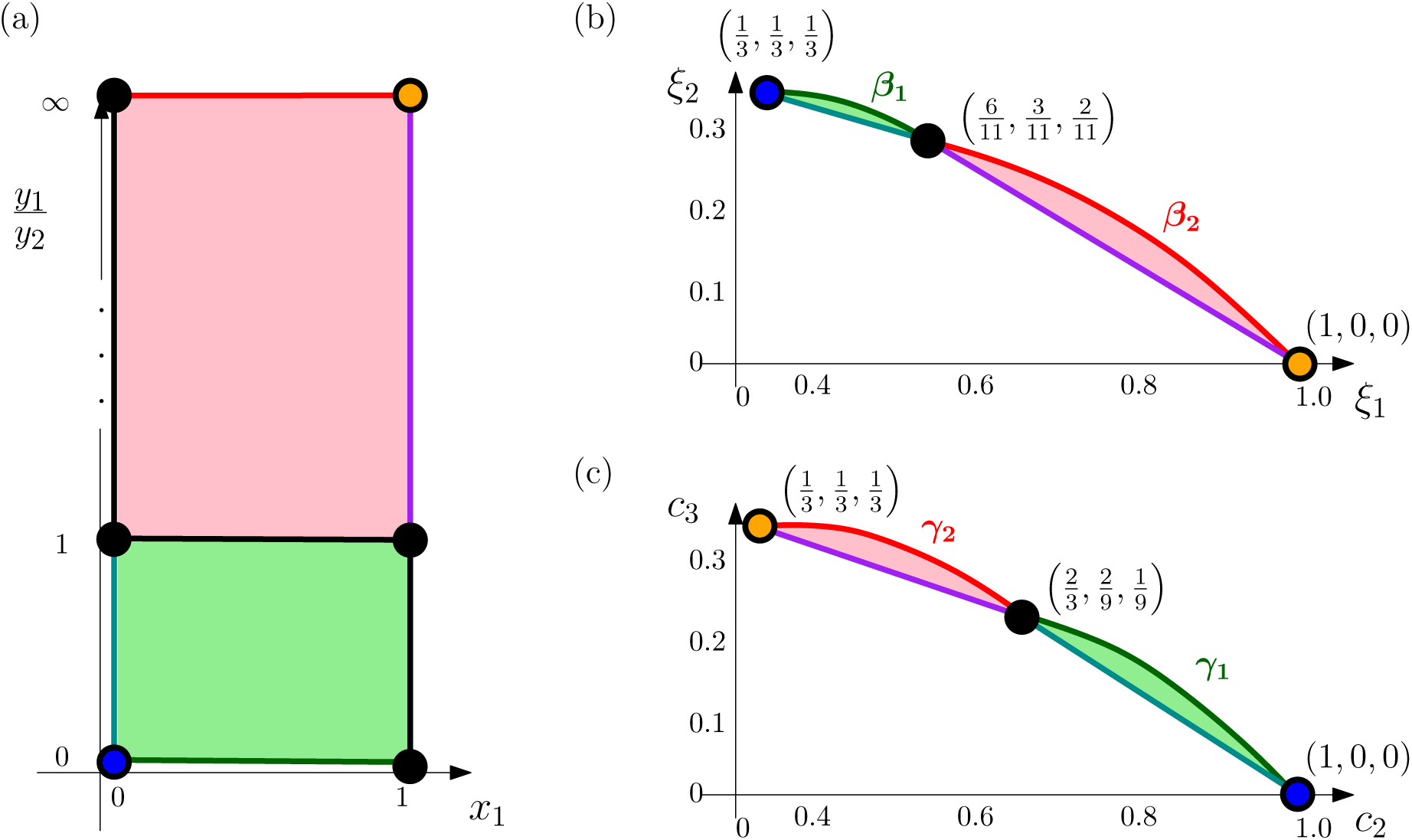
Pairing the boundaries of demography space and 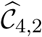. (a) The domain of χ_4,2_. Note that for fixed *y*_1_/*y_2_*, the normalized coalescence vector is the same. (b) The normalized SFS manifold 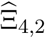 projected onto its first two coordinates. (c) The normalized coalescence manifold 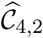 projected onto its first two coordinates. The red square at left corresponding to *y*_1_ > *y*_2_ maps to the red regions at right; the green square at left corresponding to *y*_2_ < *y*_1_ maps to the green regions at right. The black line segments on left (corresponding to *y*_1_/*y*_2_ = 1; *y*_2_ < *y*_1_ and *x*_1_ = 0 (equivalently *t*_1_ = ∞); *y*_2_ > *y*_1_ and *x*_1_ = 1 (equivalently *t*_1_ = 0)) all map to the central black points on right, since they each mimic a constant demography. The green line corresponding to *y*_1_ = 0 maps to the curve *β*_1_ in 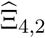 and the curve *γ*_1_ in 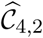; the red line corresponding to *y*_2_ = 0 maps to the curve *β*_2_ in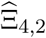 and the curve *γ*_2_ in 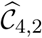. The orange point (*x*_1_ = 1,*y*_2_ = 0) maps to 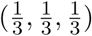 in 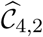 and maps to (1,0,0) in 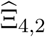. The blue point (*x*_1_ = 0,*y*_1_ = 0) maps to (1,0,0) in 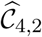 and 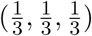 in 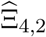. The remaining aqua and violet segments map to the segments of the same color.

We note that the map fails to be one-to-one within the domain only when *y*_1_ = 1; this is also in the pre-image of the point 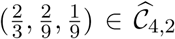. The inverse function theorem implies that on the complement of *y*_1_ = 1, the map is a homeomorphism. This is consistent with our observation that the two rectangles in Figure 3(a) correspond to the two envelopes in Figure 3(c).

### Proposition 3.3

*For all values k ≥ 3, the manifold* 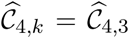, *and* 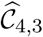 *is the interior of the convex hull of the following curve*:

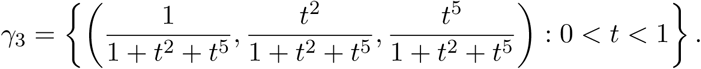

As we can see from Proposition 3.3, 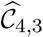 is open and convex; however, we lose one useful property of the normalized map 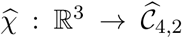. Specifically, let 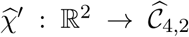 be given by 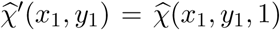, noting that 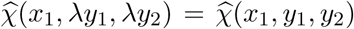 for *λ* > 0. Under this definition 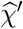 is generically one-to-one. Meanwhile, the analogous construction 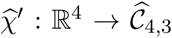 mapping the three-epoch demography with breakpoints (*x*_1_,*x*_2_) and population sizes (*y*_1_, *y*_2_,1) to the corresponding normalized coalescence vector has two-dimensional pre-images, generically. For this reason, contour images do not lend themselves to easy description. Still, we can at least describe the image of the map on the boundaries of our domain.

The easiest way to visualize the map is first to understand how the time variables affect the value of the columns of *M*_1_(4,3) and to view the *y* variables as specifying points in the convex hull of those 3 columns. The boundaries of the square (*x*_1_,*x*_2_) ∈ [0,1] × [0,1] map the columns (after rescaling to the simplex) as follows:

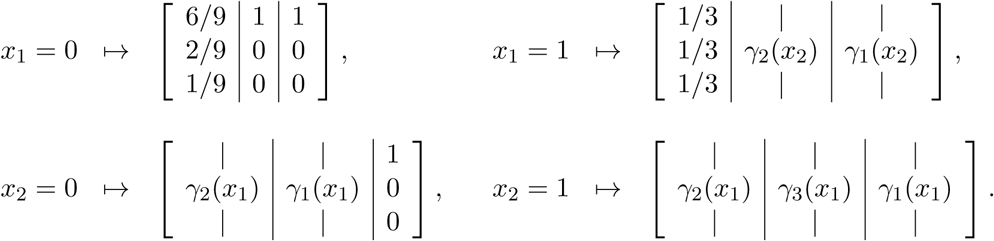

The case of *x*_2_ = 1 is the most interesting: when we fix *y*_1_ = *y*_3_ = 0 and *y*_2_ = 1, we obtain the boundary curve *γ*_3_(*t*). Note that *x*_2_ = 1 corresponds to a second epoch of length 0. The intuition is that very short population booms at the second epoch lead to coalescence vectors close to *γ*_3_. The maps encoded by a general column of *M*_1_ (4, *k*) correspond to the interior of the orange region. Adding in convex combinations of points gives the lined region, which is the remainder of C_4_,_3_; this is discussed more rigorously in Section 7. When the number of epochs *k* steps higher, all columns of *M*_1_(4, k) still map to the same region of the simplex, so 𝒞_4,*k*_ will still be contained in this convex hull. The region C4,3 is depicted in Figure 4(a).

**Figure 4:**
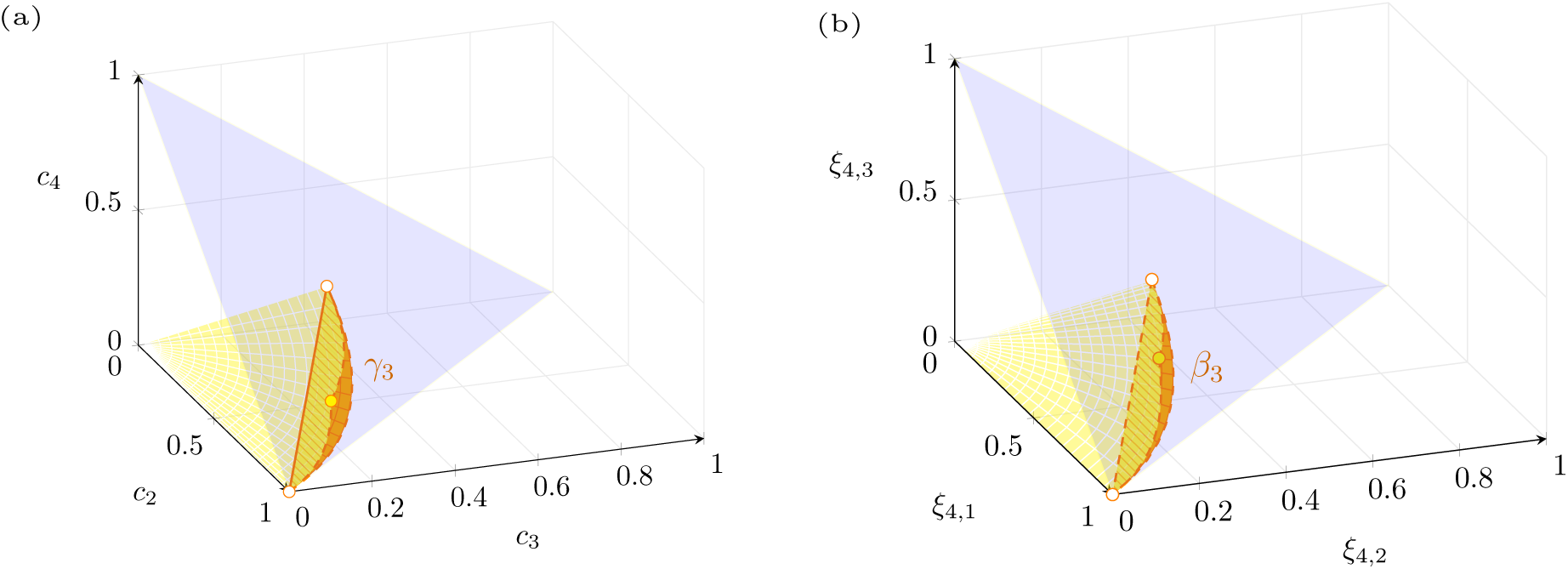
Coalescence and SFS manifolds for sample size 4 and 3 population epochs. (a) The coalescence manifold 𝓒_4,3_ is the entire yellow and orange region. The 2-simplex, shaded in blue, intersects 𝓒_4,2_ in the normalized coalescence manifold 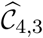. The orange region of 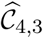, bounded by *γ*_1_, *γ*_2_, and *γ*_3_, is the image of the surface described by the columns of M_1_(4,3), while the yellow region adds in vectors gained by using linear combinations. (b) The SFS manifold Ξ_4,3_ is the entire yellow and orange region. The 2-simplex intersects Ξ_4,3_ in the normalized SFS manifold 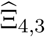. Ξ_4,2_ is obtained from 𝓒_4,3_ by a linear transformation. The orange region of 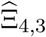, bounded by *β*_1_, *β*_2_, and *β*_3_, is the image of the surface described by the columns of *M*_1_(4,3), while the yellow region adds in vectors gained by using linear combinations.

As mentioned earlier, the SFS manifold Ξ_*n,k*_ is merely a linear transformation of *𝒞_a,k_*; however, since it is of interest in its own right, we include the formulae for Ξ_4,*k*_ analogous to those derived in this section.

### Proposition 3.4

*The following hold for the normalized* (4, *k*)-*SFS manifold*:

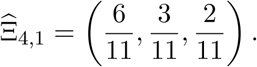

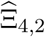*is the union of* 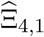 *with the convex hulls of two curves*:

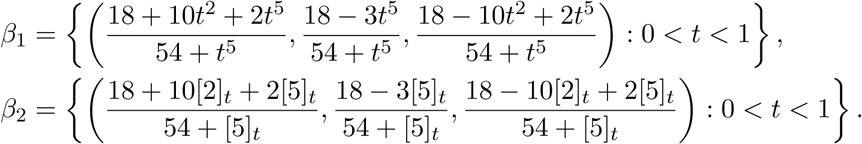

*Here, also*, [*n*]*_t_ denotes* 1 + *t* + … + *t^n^*. *Finally*, 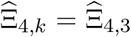 *for all k, and* 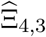 *is the convex hull of β*_3_, *where*

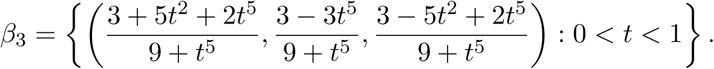

Visualizations of Ξ_4,2_ and Ξ_4,3_ may be found in Figure 1(b) and Figure 4(b).

## 4 The Ξ_*n,k*_SFS Manifold: General Properties

In this section, we examine the constant *κ_n_*, defined in Section 2 as the smallest index for which *𝒞_n,k_* ⊆ *𝒞_n,κ_n__* for all *k*. The tools for the proofs in this section come from algebraic geometry (for the derivation of the lower bound) and convex geometry (for the upper bound).

The gist of the algebraic geometry argument is that, under the *M*_2_(*n, k*) formulation, the manifold *𝒞_n,k_* can be seen to be a relatively open subset of an algebraic variety (manifold) built by a sequence of well-understood constructions. Details of this perspective are reserved for the Proofs section.

Two concrete consequences follow from this observation:

1. the ability to compute all equations satisfied by *𝒞_n,k_* using computer algebra, and
2. a formula for the dimension of the coalescence and SFS manifolds.

While the former is harder to explain without more setup, the latter can be formulated as follows:

### Proposition 4.1

*The dimension of* 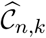 *is given by*:

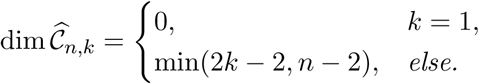

*In particular, *𝒞_n,k_* ⊊ 𝒞*_*n,k*+1_ *for* 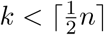.

We will illustrate how these algebraic ideas can be applied in the next case we have not seen, namely to the sample size *n* = 5.

### Example 4.2

Note that 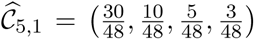, by Proposition 3.1. We will use the new ideas above to describe 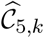 for higher values of k.

Since the normalized coalescence manifold has dimension min(2*k* − 2, *n* − 2), we know that 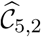 has dimension 2 inside of the 3-simplex; therefore, we anticipate that it will satisfy one equation, matching its codimension. The degree of the algebraic variety implies that this polynomial should have degree 8. Indeed, when we compute this equation using `Macaulay2` (Grayson and Stillman, 2002), we obtain a huge degree-8 polynomial with 105 terms, whose largest integer coefficient is 5, 598, 720. Finally, 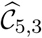 is full-dimensional in the 3-simplex, so it will satisfy no algebraic equations relative to the simplex. It would be defined instead by the inequalities determining its boundary.

While Proposition 4.1 is useful for analyzing individual coalescence manifolds, it also leads to the observation that 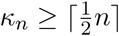, since the inclusions are proper until that index. It is worth remarking that a slightly weaker lower bound of 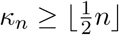 follows immediately from the identifiability result of Bhaskar and Song (2014, Corollary 7), which states that for a piecewise-constant population size function with *k* pieces, the expected SFS of a sample of size *n* ≥ 2*k* suffices to uniquely identify the function.

The convex geometry argument is more elementary. As we noted, the *M*_1_ formulation is con-tained in the convex hull over the surface described by a general column of *M*_1_. Because the columns are related, our selection of points in the surface is not unrestricted. For this reason, it is not obviously *equal* to the convex hull. However, once we fix some collection of values *x*_1_,…, *x_k_* for *𝒞_n,k_* we can use convex geometry for the resulting polytope. In particular, we use Carathéodory’s Theorem (Carathéodory (1907) or Barvinok (2002, Theorem 2.3)), which states that for *X* a subset of ℝ_*n*_ every *x* ∈ cone(*X*) can be represented as a positive combination of vectors *x*_1_,…, *x_m_* ∈ *X* for some *m* ≤ *n*.

The argument, roughly, allows us to construct any point in that convex hull, with as few as *n* + 1 points. This allows us to place the point in *𝒞_n,j_* for *j* ≤ 2*n* − 1. Since no new SFS are generated by using more than 2*n* − 1 epochs, we learn that κ_*n*_ is bounded above by 2*n* − 1.

Combining the two bounds obtained in this section, we have the following:

### Theorem 4.3

*For any integer n* ≥ 2, *there exists a positive integer κ_n_ such that* Ξ_n,k_ ⊆ Ξ_n,κ_n__ *for all k* ≥ 1. *Furthermore, κ_n_ satisfies*

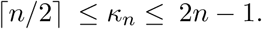

This allows us to express the SFS from any piecewise-constant demography as coming from a demography with relatively few epochs. Because the SFS is an integral over the demography, the SFS from a general measurable demography can be uniformly approximated by a piecewise-constant demography with sufficiently many epochs. Our results imply that it can be precisely obtained by a demography with at most 2*n* − 1 epochs.

## 5 Implications for Statistical Demographic Inference from Data

The SFS data used for demographic inference in population genomic studies are noisy observations of the expected SFS from the underlying population demography. Finite sequence lengths, ancestral/derived allele confounding, and sequencing and variant calling errors are some common reasons for the empirical SFS observed in sequencing studies differing substantially from the expected SFS for the underlying demographic model. It is thus possible that the empirical SFS observed in a sequencing study is not contained in the space of expected SFS Ξ_*n,κ_n_*_ for any demo-graphic model. Commonly used demographic inference methods such as *∂*`a`*∂*`i` (Gutenkunst et al., 2009), `fastsimcoal2` (Excoffier et al., 2013), and `fastNeutrino` (Bhaskar et al., 2015) perform parametric demographic inference by searching for demographies which maximize the likelihood of the observed SFS 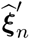. Under the widely used Poisson Random Field model which assumes that the genomic sites being analyzed are unlinked (Sawyer and Hartl, 1992), maximizing the likelihood is equivalent to minimizing the KL divergence between the empirical SFS and the expected SFS under the parametric demographic model. Given an observed SFS 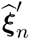, these algorithms traverse the interior of some user-specified space of parametric population size functions such as Π_*k*_, while computing the expected SFS 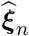 under the forward map 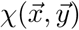 in each optimization iteration. The optimization procedure either terminates and returns a demography *η* in the search space Π_*k*_ whose expected SFS 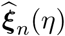 minimizes the KL divergence KL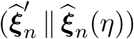 among all demographies in Π_*k*_, or it exhibits runaway behavior where some population sizes or epochs diverge to infinity or go to 0 with successive optimization iterations.

Our geometric study of the 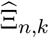 SFS manifold can clearly explain the success and failure modes of these optimization algorithms. Suppose the demographic search space is Π_*k*_. When the observed SFS 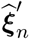 lies in the interior of the 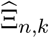 SFS manifold, this observed SFS is also exactly equal to the expected SFS of some demographic model *η*^*^ ∈ Π_*k*_, and hence any of the above mentioned optimization algorithms, barring numerical difficulties,^†^ should be able to find this demography *η*^*^ whose expected SFS 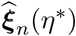 is exactly equal to the observed data 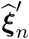 and KL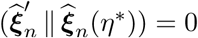. On the other hand, if the noise in the observed SFS causes it to lie outside the 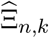 SFS manifold, these optimization algorithms will attempt to find the demography η(*t*) ∈ Π_*k*_ which minimizes the projection under the KL divergence of the observed SFS 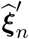 onto the 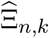 SFS manifold,

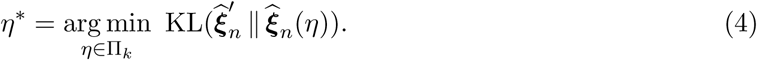

The optimization problem in (4) has a couple of issues. As described in Sections 3 and 4, the SFS manifold 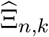 is a set where most of the boundary points are not contained in 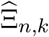. For example, the description of 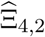 in Section 3 showed that the only boundary point of 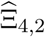 contained in 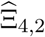 is the expected SFS corresponding to the constant population size function, namely the point (6/11, 3/11, 2/11). Since the points on the boundary of the 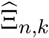 correspond to different limiting regimes with the epoch durations and population sizes tending to 0 or ∞, commonly used demographic inference algorithms that attempt to solve the optimization problem in (4) experience runaway behavior when the observed SFS lies outside the 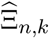 SFS manifold. Figure 5 shows the SFS vectors of simulated sequences (blue circles) under the coalescent with a constant population size, where most of the simulated SFS fall outside the 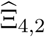 manifold. We used `fastNeutrino` to fit two-epoch piecewise-constant demographies to these simulated SFS. The observed SFS vectors which project onto the curved boundary of the upper convex set are inferred to come from a demography where both the population size and the duration of the recent epoch, *y*_1_ and *t*_1_ respectively, go to 0, while for the observed SFS projecting onto the curved boundary of the lower convex set, *y*_1_ and *t*_1_ diverge to infinity. In both cases, the location of the projection along the curved boundaries is determined by the value of *x*_1_ = exp(−*t*_1_/*y*_1_) where *x*_1_ ∈ (0,1). On the other hand, for the observed SFS which project onto the straight line boundaries of the upper and lower convex sets, the inferred recent epoch durations go to 0 (upper convex set) or diverge to infinity (lower convex set) while the inferred recent population size *y*_1_ is a constant relative to the ancient population size *y*_2_ and the location of the projection along the boundaries is determined by *y*_1_/*y*_2_.

**Figure 5:**
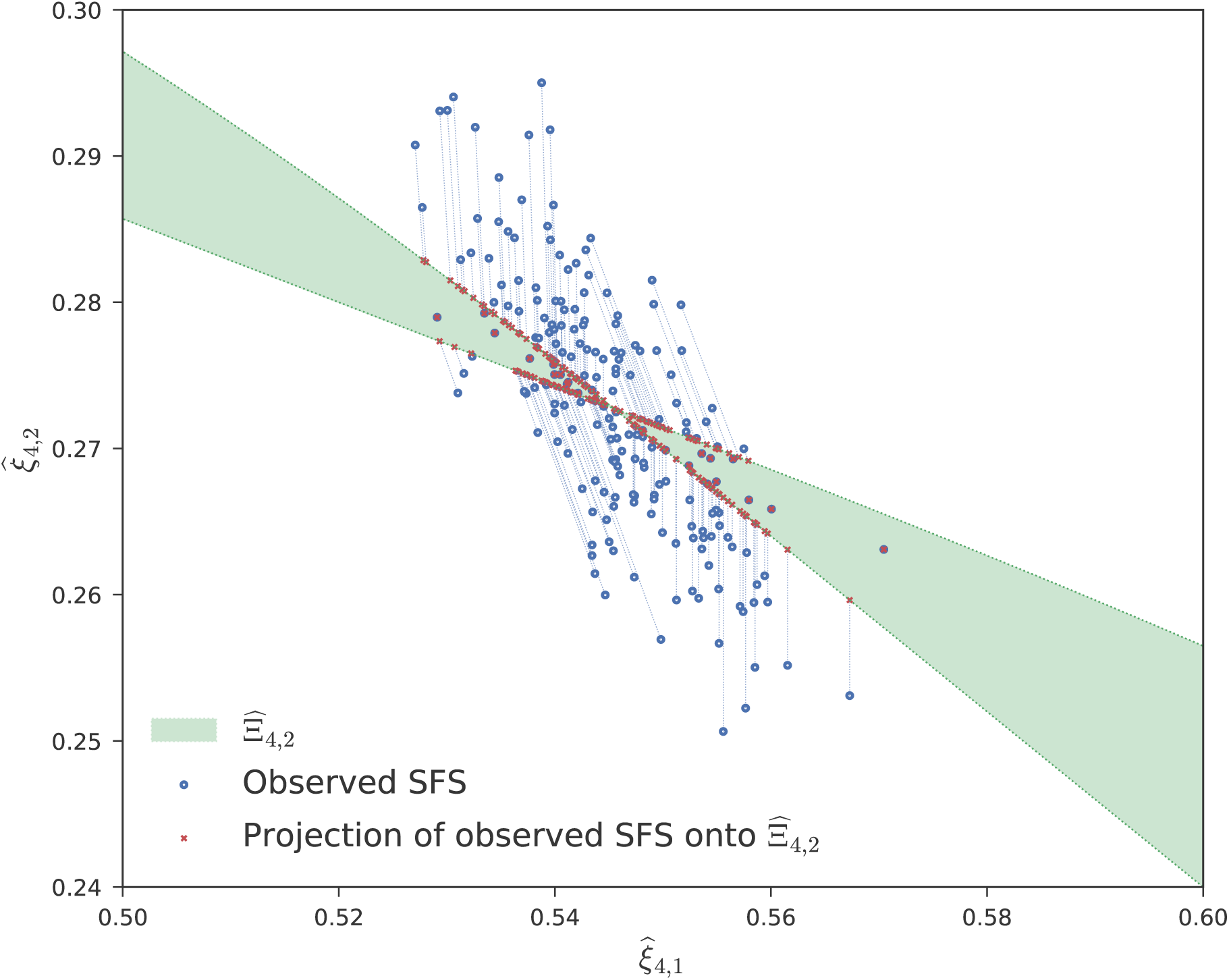
Each blue circle is the observed SFS of *n* = 4 haplotypes simulated using msprime (Kelleher et al., 2016) under a constant population size coalescent with recombination using realistic mutation and recombination rates of 10^−8^ mutations and 2.2 × 10^−8^ crossovers per basepair per generation per haploid. Each sequence has 1000 unlinked loci of length 10 kb each, resulting in an average of 7,300 segregating sites. The red crosses are the expected SFS at the two-epoch piecewise-constant demographies inferred for these simulated SFS using fastNeutrino; the red crosses are the projections of the observed SFS onto the closure of 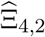 using the KL divergence, with the dotted blue lines showing the correspondence between the observed SFS and their projections. For observed SFS lying in the interior of 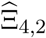, the observed SFS and their projections coincide, while the observed SFS lying outside 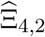 project onto the boundaries of one of the two convex sets that form 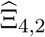.

A second more subtle issue arises from the fact that the 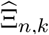 SFS manifold being projected onto in (4) may be a non-convex set, and hence the solution to the optimization problem in (4) may not be unique.^‡^ For example, we already observed in Section 3 that the set 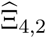 is non-convex and is given by the union of two convex sets. Hence, for *n* = 4 and *k* = 2, and for some values of the observed SFS 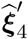, there could exist multiple different demographic models η_1_,η_2_ ∈ Π_2_, η_1_ ≠ η_2_, such that

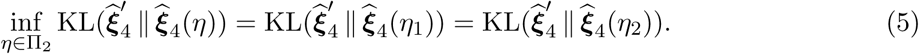

By algebraic considerations, the observed SFS 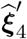 which have such non-unique projections onto 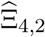 form a set of measure zero among all possible probability vectors on three elements, and hence such SFS are unlikely to be encountered in real data. However, the existence of such SFS vectors 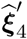 with non-unique projections implies that slight perturbations to these vectors, say due to different quality control procedures for selecting the set of genomic sites to analyze, could result in very different demographic models being inferred due to the projection of the perturbed vector occurring onto one or the other of the two convex sets composing 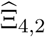. This is also apparent in Figure 5, where several pairs of observed SFS vectors that are very close to each other project onto the different convex sets forming 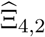. As described in the previous paragraph, the boundaries of the upper and lower convex sets represent various limiting regimes with either vanishingly small or arbitrarily large recent population sizes and epoch durations, and this shows that even minor perturbations to the SFS vector could yield qualitatively very different inference results which cannot be reliably interpreted.

## 6 Discussion

In this work, we characterized the manifold of expected SFS *Ξ_n,k_* generated by piecewise-constant population histories with k epochs, while giving a complete geometric description of this manifold for the sample size *n* = 4 and *k* = 2 epochs. This special case is already rich enough to shed light on the issues that practitioners can face when inferring population demographies from SFS data using popular software programs. While we demonstrated these issues in Section 5 using the `fastNeutrino` program, the issues we point out are *inherent* to the geometry of the SFS manifold and not specific to any particular demographic inference software. Our simulations showed that the demographic inference problem from SFS data can be fraught with interpretability issues, due to the sensitivity of the inferred demographies to small changes in the observed SFS data. These results can also be viewed as complementary to recent pessimistic minimax bounds on the number of segregating sites required to reliably infer ancient population size histories (Terhorst and Song, 2015).

Our investigation of piecewise-constant population histories also let us show a general result that the expected SFS for a sample of size *n* under *any population history* can also be generated by a piecewise-constant population history with at most 2*n* − 1 epochs. This result could have potential applications for developing non-parametric statistical tests of neutrality. Most existing tests of neutrality using classical population genetic statistics such as Tajima’s *D* (Tajima, 1989) implicitly test the null hypothesis of selective neutrality *and* a constant effective population size (Stajich and Hahn, 2004). Exploiting our result characterizing the expected SFS of samples of size *n* under arbitrary population histories in terms of the expected SFS under piecewise-constant population histories with at most *κ_n_* epochs, we see that the KL divergence of an observed SFS 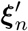 to the expected SFS **ξ**_*n*_(η*) under the best fitting piecewise constant population history *η*^*^; ∈ *Π_η_n__* with at most *κ_n_* ≤ 2*n* − 1 epochs is also equal (up to a constant shift) to the negative log-likelihood of the observed SFS 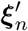 under the best fitting population size history without any constraints on its form, assuming the commonly used Poisson Random Field model where the sites being analyzed are unlinked. One can then use the KL divergence inferred by existing parametric demographic inference programs to create rejection regions for the null hypothesis of selective neutrality without having to make any parametric assumption on the underlying demography. Such an approach would also obviate the need for interpreting the inferred demography itself, since the space of piecewise-constant population histories is only being used to compute the best possible log-likelihood under any single population demographic model. This approach could serve as an alternative to recent works which first estimate a parametric demography using genome-wide sites, and then perform a hypothesis test in each genomic region using simulated distributions of SFS statistics like Tajima’s *D* under the inferred demography (Rafajlovic et al., 2014). We leave the exploration of such tests for future work.

## 7 Proofs

### Proof of Proposition 2.1

First, we reduce the integral expression for *c_m_* to a finite sum; then we make appropriate manipulations until we arrive at the desired expressions.

Coalescence in the Wright-Fisher model is an inhomogeneous Poisson process with parameter 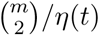. Therefore, the probability density of first coalescence at time *T* is:

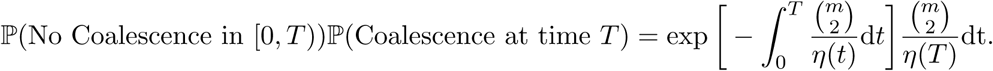

Let 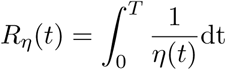. To compute the expected time to first coalescence, we have the integral:

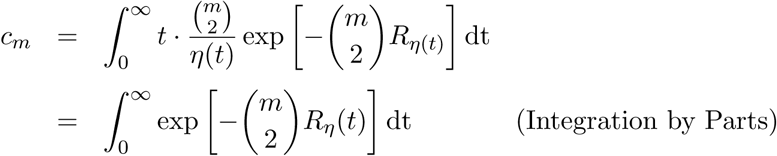

Substituting variables, *τ* = *R_η_*(*t*), note that d*τ* = *η*(*R*^−1^(*τ*))d*τ*. Therefore, the integral becomes:

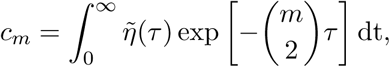

where 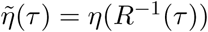.

The population size *η*(*t*) is a piecewise constant function, whose value *η*(*t*) = *η_j_* if *t*_*j*-1_ ≤ *t* ≤ *t_j_*. As specified in the Proposition, *t*_0_ = 0, *t_k_* = ∞, and (*y*_1_,…,*y_k_*) is the vector of population sizes. Observe that 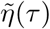 is also piecewise constant. In particular,

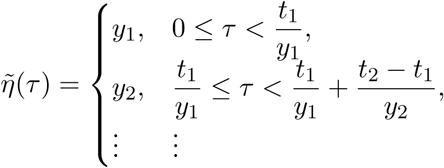

Let *s_j_* = *t_j_* − *t*_*j*-1_ for brevity. The resulting formula is:

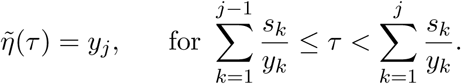

We turn the integral into a sum of integrals on the constant epochs:

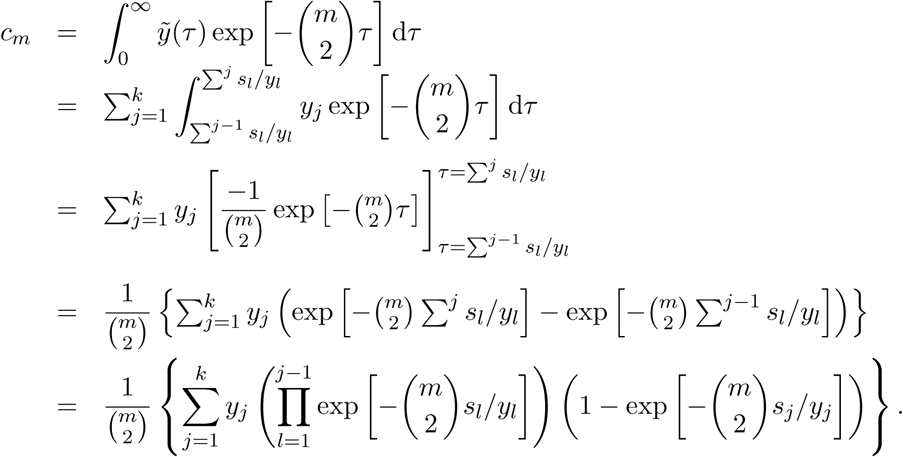

We now make the substitution *x_j_* = exp [–*s_j_*/*y_j_*]. Note that the old restriction *t*_*j*+1_ > *t_j_* > 0 becomes the new constraint 0 < *x_j_* < 1. Our formula for the *c_m_* is now:

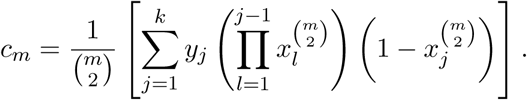

Noting the linear form of this expression, we factor as a matrix multiplication:

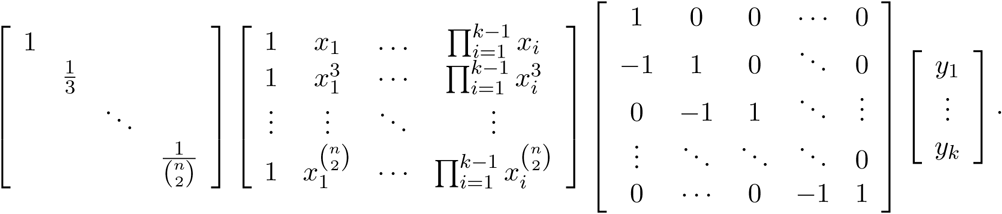

Combining the first three matrices yields (2); combining the first two and last two separately yields (3).

### Proof of Proposition 3.1

We justify each equation in turn:

1. As mentioned in the introduction, this is a classical result in population genetics, and can be derived directly from (3).
2. The inclusion 𝒞_2,1_ ⊂ 𝒞_2,*k*_ is immediate, so we need only show that any *a* ∈ 𝒞_2,*k*_ satisfies *a* > 0. Using (2), a is written as a sum of products of strictly positive numbers; so 𝒞_2,*k*_ ⊂ 𝒞_2,1_.
3. First, we show that 𝒞_3,2_ is the interior of the open cone spanned by (1, 0) and (1,1). Fix *y*_1_ = *a*/(1 — *x*_1_) (for a positive) and consider χ(*x*_1_,*a*/(1 − *x*_1_),*y*_2_):

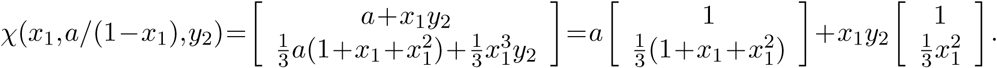 When *x*_1_ → 0, the second vector approaches (1,0); when *x*_1_ → 1, the first vector approaches (1,1). The vectors are in the interior of that cone for all other permissible values of x_1_ and y_2_. To show that 𝒞_3,*k*_ =𝒞_2,1_, note that for larger values of *k*, the same cone of vectors are produced. In particular, χ(*x*_1_,…,*x*_*k*-1_,*y*_1_,…,*y_k_*) yields

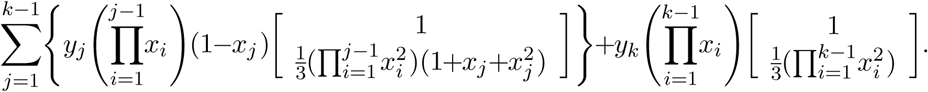 Clearly, the second coordinate of all vectors is bounded between 0 and 1.

### Proof of Proposition 3.2

First we observe that *γ*_1_ and *γ*_2_ are normalizations of the curves defined by parameterizations 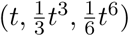 and 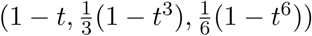 where *t* is constrained to the open interval (0,1).

Now we claim that the definition in terms of the map χ(*x, y*) is equivalent to the definition in terms of these two curves. We can use the first formulation of χ to prove this:

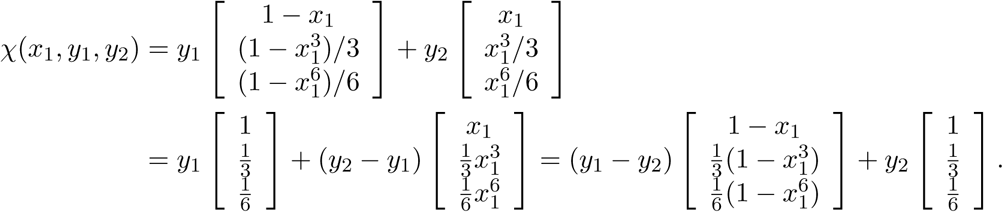

When *y*_2_ = *y*_1_, the image is the point (2/3,2/9,1/9) = *X* as stated. When *y*_2_ > *y*_1_, we can use the left-hand expression to view the image as a point on the line segment between 𝒞_4,1_ and the curve (*t*, *t*^3^/3, *t*^6^/6). When *y*_2_ < *y*_1_, the right-hand expression can be used to write the image as a point on the line segment between *X* and (1 − *t*, (1 −*t*^3^)/3, (1 −*t*^6^)/6). This means that the image of χ is contained in the regions and point specified.

To show that the reverse inclusion holds, we fix a point *P* in the interior of the convex hull of γ_1_. By convexity, the line segment from *X* to *P* is contained in the region; continue in the direction *P* − *X* until the line intersects the curve. This must occur because all points in the region are further from the bounding line than *X*. The point of intersection *q* is specified as *q* = γ_1_(τ) for some τ ∈ (0,1). By convexity, there exists some *ρ* such that *ρ* 𝒞_4,1_ + (1 − *ρ*)*q* = *P*. Fixing *x*_1_ = τ, *y*_1_ = *ρ* and *y*_2_ = 1, shows that *P* is in the image of *χ*. The same argument holds with slight variation for γ_2_.

### Proof of Proposition 3.3

The strategy to prove the equality of 𝒞_4,3_ and the cone over {*t*, *t*^3^, *t*^6^} comes in two steps:

1. Show that the columns of *M*_1_(4,*k*) are always contained in the region *R* whose boundary is γ_1_ ∪ γ_2_ ∪ γ_3_.
2. Divide the convex hull of R into two regions and show that each of these regions are included in 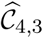.

First we demonstrate that the regions maps precisely into *R*. We have already shown in the main text of the document that the boundaries of (0,1) × (0,1) map to the boundaries of *R* under the mapping defined by 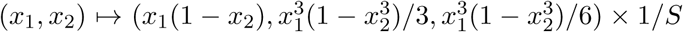, where *S* is the sum of the coordinates. We compute the Jacobian of this map explicitly in `Macaulay2` (Grayson and Stillman, 2002). The result is:

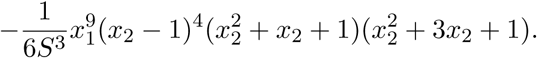

Plainly, this is nowhere zero in our domain. The inverse function theorem then implies that the interior is contained in the image of the boundaries. This accomplishes Step 1 of our proof.

For Step 2, we divide the image into two regions:

1. The triangle defined by vertices (1, 0, 0), (2/3, 2/9,1/9) and (1/3,1/3,1/3), including the two edges [(1/3,1/3,1/3), (2/3, 2/9,1/9)] and [(2/3, 2/9,1/9), (1, 0, 0)].
2. The remainder of the convex hull of *R*–explicitly, the interior of the region bounded by γ_3_ and the line segment [(1/3,1/3,1/3), (1, 0, 0)].

To show that the triangle is included, let *x*_2_ = *є* ≈ 0, and let *x*_1_ vary. Then the third column sits arbitrarily close to (1,0,0) and the first column traces out γ_2_. Set *y*_2_ ≈ 0 and toggle *y*_1_ and *y*_3_, to obtain the full span, including the interior of the triangle, and the line segment [(1/3,1/3,1/3), (2/3, 2/9,1/9)]. Set *x*_1_ = 1 − *є*, and the first column sits at (1/3,1/3,1/3) while the third column traces out γ_1_. This catches the missing line segment.

For the remainder of the convex hull, fix a point P in this region. This point lies on a line segment between (2/3,2/9,1/9) and some point *Q* in γ_3_. Suppose it is equal to *ρ* · (2/3, 2/9,1/9) + (1 − *ρ*) · *Q*. Set *x*_2_ = 1 − *є* ≈ 1. We can choose *є* and *x*_1_ so that the second column is arbitrarily close to P. Furthermore, observe that the first column is approximately equal to the point on *γ*_2_ corresponding to x_1_ and the third column is approximately the point on *γ*_1_ corresponding to *x*_1_. Choosing *y*_1_ = *y*_3_ = *ρ* and *y*_2_ = 1 − *ρ* points us to

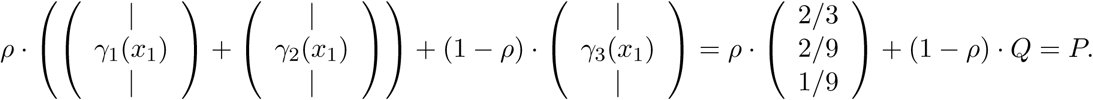

### Proof of Proposition 3.4

This is a direct application of the linear map *W*_4_, computed as in Polanski and Kimmel (2003):

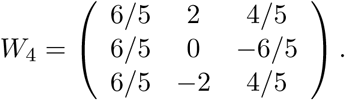

**Proof of Proposition 4.1**

In order to prove the result about dimension, we show that 𝒞_*n,k*_ is a relatively open subset of a certain algebraic variety. Because the relevant operations are native to projective geometry, we transport our objects of interest in the obvious way to projective space. The same scaling properties that allow us to focus on the simplex also lead to good behavior in projective space.

**Lemma 7.1.** *For k ≥* 2, *the Zariski closure of* 𝒞_*n,k*_ *is the affine cone over 𝓘(*σ*_*k*-2_(*C_n_*,*p_n_*)), where*:

1. *C_n_ is the projective curve defined by mapping* [*s : t*] *to*

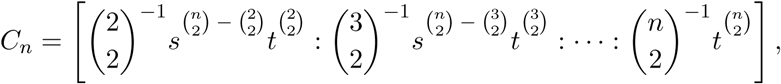
2. *p_n_ is the projective point* 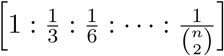,
3. *𝓘 denotes the join of algebraic varieties, and*
4. *σ_i_(·) denotes the i-th secant variety. Following Harris (2013), the i-th secant variety is the union of i-dimensional planes generated by i* + 1 *points in the variety*.

*Proof of Lemma 7.1.* The variety *ϕ*(*σ*_*k*-2_(*C_n_*),*p_n_*)) is the image of the following map:

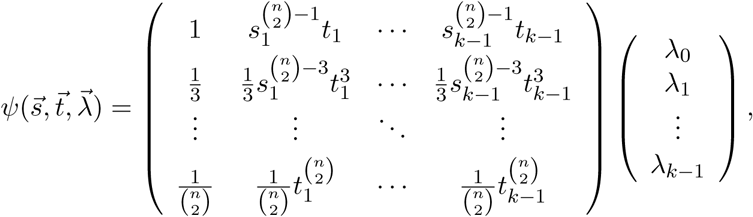

where *s_i_* and *t_i_* are not simultaneously zero, and λ is unrestricted.

Define the map *ϕ* : ℝ^2*k*-1^ → (ℙ^1^)^*k*−1^ × ℝ^*k*^ sending

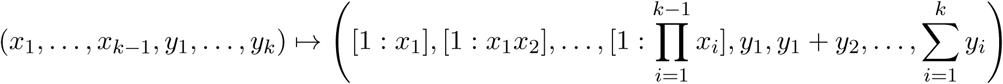

We can recast the expression in (3) as the composition *ψ* o *ϕ*. Based on this formulation, the set 𝒞_*n,k*_ is clearly contained in 𝓘(*σ*_*k*-2_(*C_n_*),*p*). To demonstrate the equality of the Zariski closures, we only need to show that the dimensions match and that the variety is irreducible. Both joins and secants have the property that irreducible inputs yield irreducible outputs, so the variety of interest is irreducible. The image of *ϕ* is open in (ℙ^1^)^*k*-1^ × (ℙ^1^)^*k*−2^, and the map *ψ* has deficient rank on a set of positive codimension. Therefore, the composition of *ψ* o *ϕ* has full dimension. This proves the Lemma.

The *i*-th secant variety of an irreducible nondegenerate curve in ℙ^n^ has projective dimension given by min(2*i* + 1,*n*) (Harris, 2013, Exercise 16.16). The curve *C_n_* is a toric transformation of a coordinate projection of the rational normal curve. The rational normal curve is nondegenerate, and both of these operations preserve that property. This means our secant variety has projective dimension min(2(*k* − 2) + 1,*n* − 2) = min(2*k* − 3,*n* − 2). The join with a point adds 1 to the dimension of the variety, while the operation of passing to the affine cone adds 1 to the dimension of the variety and the ambient space. However, normalizing to the (*n* − 2)-simplex subtracts 1 from both variety and ambient space again. This means that dim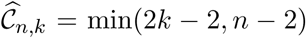, assuming that *k* ≥ 2.

### Proof of upper bound in Theorem 4.3

Suppose a point c is in *𝒞_n,q_*. By definition, this implies that there is a point (*x*_1_,…, *x*_*q*-1_, *y*_1_,…, *y_q_*) such that (2) yields

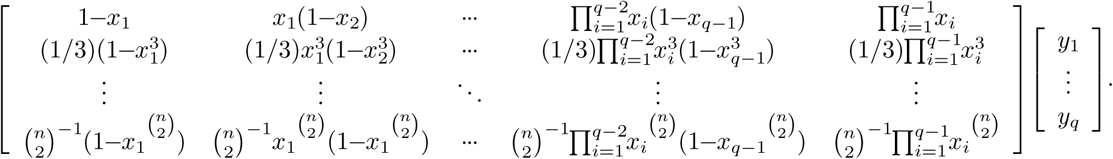

Since the point c is in the cone over the *q* columns of the matrix, Carathéodory’s Theorem implies that it is also in the cone over some *n* − 1 of the columns. Therefore we can replace the vector *y*_1_,…, *y_q_* with 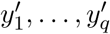 so that all but *n* − 1 (or fewer) are zero.

Passing to the expression in (3), this gives us:

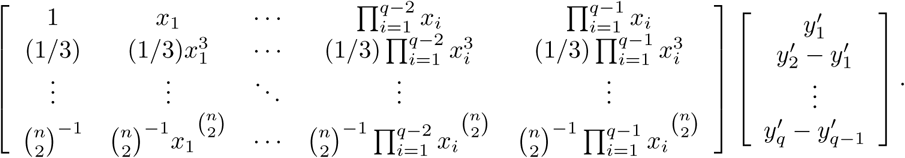

Since at most *n* − 1 of the 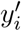 are nonzero, at most 2*n* − 2 of the indices of the vector at right are nonzero. We can delete the columns of the X matrix corresponding to zero entries except the first column. A new sequence 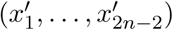 can then be obtained from the ratio between the first entries in adjacent columns. The new sequence 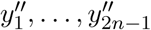 can be obtained by taking the sequence of partial sums of the vector.

## Acknowledgments

We thank the Simons Institute for the Theory of Computing, where some of this work was carried out while the authors were participating in the “Evolutionary Biology and the Theory of Computing” program. This research is supported in part by a Math+X Research Grant, an NSF grant DMS-1149312 (CAREER), an NIH grant R01-GM109454, and a Packard Fellowship for Science and Engineering. YSS is a Chan Zuckerberg Biohub investigator.

## Footnotes

* The sets *Ξ_n,k_* and 𝒞_*n,k*_ are not technically manifolds; they would be more accurately described as semialgebraic sets. However, for expository purposes, we use the widely known term “manifold”.

† The package ∂`a`∂`i` uses numerical methods to approximate the solution to a diffiusion PDE, while `fastsimcoal2` uses coalescent simulations to estimate the expected SFS for a given demographic model. Hence, these software packages might have numerical issues beyond the failure modes we consider here. For this reason, we conduct our inference experiments in this section using the fastNeutrino package, which uses the analytic results of Polanski and Kimmel (2003) for exact computation of the expected SFS for piecewise-constant population size functions.

‡ If 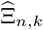 is a convex set, the solution to (4) is unique due to the fact that the KL divergence is a convex function of either argument. Namely, for 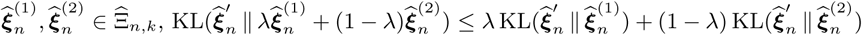 with equality holding if and only if 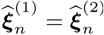.

